# Cryopreservation of Infectious *Cryptosporidium parvum* Oocysts

**DOI:** 10.1101/336040

**Authors:** Justyna J. Jaskiewicz, Rebecca D. Sandlin, Anisa A. Swei, Giovanni Widmer, Mehmet Toner, Saul Tzipori

## Abstract

Cryptosporidiosis in an enteric infection caused by *Cryptosporidium* parasites and is a major cause of acute infant diarrhea in the developing world. A major bottleneck to research progress is the lack of methods to cryopreserve *Cryptosporidium* oocysts, thus requiring routine propagation in laboratory animals. Here, we report a method to cryopreserve *C. parvum* oocysts by ultra-fast cooling. Cryopreserved oocysts exhibits high viability and robust *in vitro* excystation, and are infectious to interferon-γ knockout mice. The course of the infection is comparable to what we observe with unfrozen oocysts. Oocyst viability and infectivity is not visibly changed after several weeks of cryogenic storage. Cryopreservation will facilitate the sharing of oocysts from well characterized isolates and transgenic strains among different laboratories.

## Introduction

*Cryptosporidium* is second only to *Rotavirus* among the enteric pathogens that cause moderate-to-severe diarrhea in children aged less than two years in developing countries (*1*). In this setting, cryptosporidiosis is associated with mortality among infants aged 12-23 months, irrespective of their HIV status. Protracted and often severe diarrhea may lead to stunted growth and cognitive impairment. Cryptosporidiosis is also an opportunistic infection among immunocompromised individuals and may cause life-threatening persistent diarrhea and wasting (*2*). Current anti-parasitic drugs are ineffective in immunocompromised and malnourished individuals, creating a need for improved therapeutics. The lack of cryopreservation methods and the short shelf life of oocysts make it difficult to generate and distribute genetically characterized oocysts. Consequently, laboratory strains of *Cryptosporidium* must be maintained by propagation in susceptible animals (mice, calves and piglets) every 6-8 weeks, an expensive and time-consuming process. This limitation has precluded the sharing of well-characterized isolates among different laboratories, limiting the evaluation of therapeutics and vaccines. Despite numerous attempts, methods to cryopreserve *Cryptosporidium* have eluded scientists for many years (*3-6*). Development of cryopreservation methods for *Cryptosporidium* would thus be transformative to drug discovery efforts by enabling broader access to parasites for research and significantly, for human challenge studies to evaluate drug efficacy. Ideally, inocula for human challenge studies should be of the same optimized, standardized and validated batch of oocysts. This is currently impossible as oocysts must be produced monthly, over the two-year trial period and undergo optimization, standardization and validation *in vitro* and *in vivo* following each propagation prior to evaluation at different testing sites. The ability to cryopreserve aliquots of oocysts from a single batch and thaw as needed can ascertain uniformity of the challenge dose and thus reduce considerably batch-to-batch variation.

Two standard approaches to cryopreservation include slow cooling and vitrification. In either method, intracellular cryoprotective agents (CPAs) are typically required for successful cryopreservation. For slow cooling approaches, cells are typically treated with ∼1-2 M CPAs and cooled at a rate of ∼1°C/min until the sample is frozen (*7, 8*). In contrast, vitrification utilizes high concentrations of CPAs (∼4-8 M) combined with rapid cooling rates to achieve formation of a glassy, amorphous solid resulting in ‘ice-free` cryopreservation (*9, 10*). Compared to slow cooling, vitrification often leads to increased cell viability and function, but can be limited in use by CPA toxicity and cumbersome methodology. To address the limitation imposed by CPA toxicity, devices that enable ultra-fast cooling rates have been developed. Since the CPA concentration necessary to vitrify is inversely related to the cooling rate, these devices enable vitrification with significantly less CPA (*11-16*). Because previously reported attempts to cryopreserve *C. parvum* using slow cooling rates resulted in noninfectious oocysts (*3-6*), ultra-fast cooling was explored as a novel approach to cryopreservation.

Here, we describe a protocol to successfully cryopreserve infectious *C. parvum* oocysts using ultra-fast cooling rates of ∼4000 K/s. We first demonstrate that hypochlorite treatment is an effective method to permeabilize oocysts and enable intracellular uptake of CPAs. Subsequent characterization of CPA toxicity enables the selection of a cocktail solution containing DMSO and trehalose at concentrations that permit extracellular vitrification. To achieve ultra-fast cooling rates, oocysts are loaded into highly conductive silica microcapillaries (200 µm inner diameter) that are subsequently plunged into liquid nitrogen, such that the extracellular solution vitrifies nearly instantaneously. Using this method, thawed oocysts are viable and appear morphologically normal. Inoculation of thawed oocysts into interferon-γ (IFN-γ) knockout mice demonstrates that infectivity is maintained. This cryopreservation protocol overcomes a major bottleneck in the study of *Cryptosporidium* biology and the development of therapeutics.

## Results

### Hypochlorite treatment permeabilizes oocysts to CPA

CPA uptake by oocysts may be hindered by the highly impermeable, tetralaminar wall that protects the sporozoites from environmental exposure (*17*). Indeed, Figure 1a shows that exposure to an expectedly toxic concentration of 40% CPA for up to two hours results in negligible oocyst mortality, suggesting that little or no CPA permeation occurs. This observation was a major concern as typically intracellular presence of CPAs is required for successful cryopreservation. To increase oocyst permeability, hypochlorite treatment was explored, an approach previously demonstrated by Mazur *et al.* to increase permeability of *Drosophila* larvae (*18*). Hypochlorite is non-toxic to oocysts and bleach is often used to sterilize their surface (Supplementary Fig. 1). Figure 1b shows that following hypochlorite treatment, oocysts incubated with 40% DMSO for 30 min exhibit 70.1±6.3% toxicity compared to controls of unbleached oocysts where no toxicity is observed. Additionally, Supplementary Fig. 2 indicates that bleached oocysts initiate uptake of CPA after >3 min of incubation. While these data indirectly suggest that CPA permeability is achieved, it is not known what fraction of CPA actually accumulates within the oocyst. Insufficient uptake of CPA would result in a high intracellular water content, leading to the potential formation of lethal intracellular ice crystals upon rapid cooling. In order to reduce intracellular water content, we characterized hyperosmotic solutions of NaCl and trehalose, a non-penetrating, extracellular CPA, as a method to dehydrate oocysts. Figure 1c shows that incubation in increasing concentrations of trehalose leads to decreasing oocyst volumes, consistent with dehydration. Interestingly, NaCl did not have a significant effect on oocyst volume, although it is commonly used to exclude water from cells. Despite severe dehydration (up to 71±8.4% volume reduction) in hyperosmotic solutions of trehalose, only marginal toxicity was observed in rehydrated oocysts (Figure 1d).

**Fig 1:**
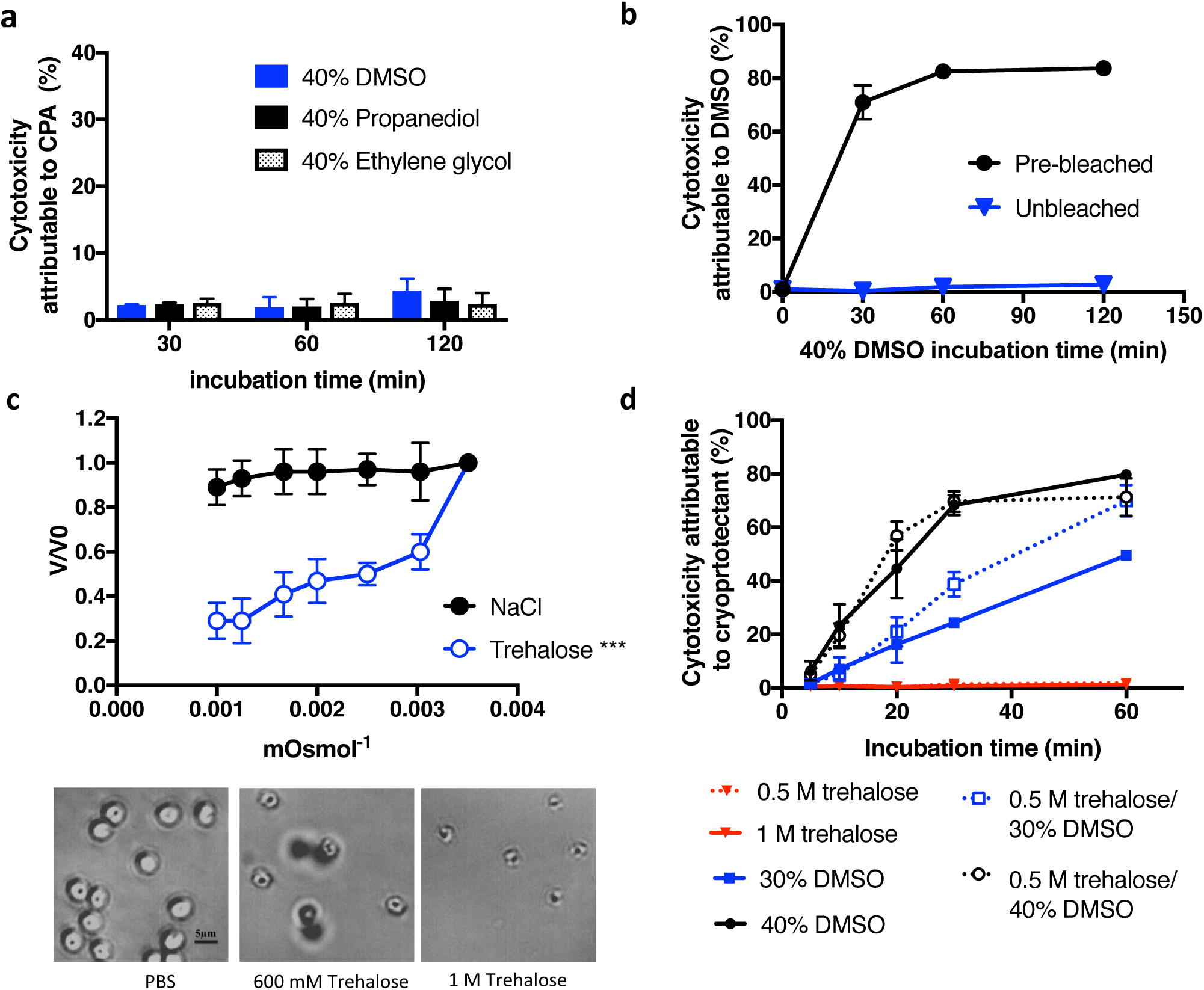
*C. parvum* oocyst response to CPAs. **a** Oocysts were incubated with high concentrations of several common CPAs. As no significant increase in oocysts toxicity was observed over long incubation intervals (30-120 min), it was concluded that the oocysts are impermeable to CPAs (n=3). **b** Hypochlorite treatment was used as a method to increase oocyst permeability to CPAs. Bleached and unbleached oocysts were incubated with 40% DMSO and cytotoxicity was quantified by flow cytometry using PI exclusion (n=3). Both bleached and unbleached oocysts incubated with PBS served as a normalizing control. While unbleached oocysts exhibited no toxicity, bleached oocysts incubated with DMSO exhibited substantial toxicity over the observed period (30-120 min), suggesting CPA permeability was achieved. **c** Oocyst volume was measured by flow cytometry to characterize dehydration induced by hyperosmotic solutions of NaCl and trehalose (V, estimated oocyst volume; Vo, volume of control oocysts). While NaCl had insignificant effect on volume (one-way ANOVA; *p*=0.90, *f*=0.33, *df*=20), significant dehydration was observed using trehalose (one-way ANOVA; *p*<0.0001, *f*=18.65, *df*=20), a common extracellular CPA. Data met requirements of normality (Wilk Shapiro test; *p*>0.29 and *p*>0.85 for all NaCl and trehalose concentrations, respectively) and homoscedasticity (Brown-Forsythe test; *p*=0.80 and *p*=0.88 for NaCl and trehalose, respectively) for inclusion in ANOVA analysis. Panels show a significant decrease in cell volume in response to increasing concentrations of trehalose compared to the untreated PBS control (n=3). Micrographs show oocysts exposed to trehalose or PBS as indicated. **d** The kinetics of CPA toxicity were measured by incubating bleached (5% bleach, 1 min) oocysts in DMSO, with our without dehydration in trehalose, for 5-60 min. Oocysts were dehydrated with 1 M trehalose for 10 min and then treated with a solution of DMSO to achieve a final concentration of 0.5 M trehalose/30-40% DMSO, or 30-40% DMSO only, as measured by PI inclusion (n=3). All values indicate means and error bars indicate standard deviation.

### Kinetics of CPA toxicity and role of oocyst age

Due to the unique biophysical and biochemical properties among oocyst stocks, the optimal CPA identity, concentration and incubation period that leads to maximal loading and minimum cytotoxicity was determined. Figure 1d shows the kinetics of toxicity of bleached oocysts incubated with various concentrations of DMSO, trehalose or both. The results indicate that following a 20 min incubation in 30% DMSO, with or without trehalose, ∼20% (20.9±5.5% with and 16.3±6.8% without trehalose) cytotoxicity is observed. This suggests that CPA permeation occurs, but at a concentration that does not lead to high toxicity. It became apparent that significant variation in CPA toxicity was present between oocyst stocks of different ages, presumably related to structure and permeability of the oocyst wall (Supplementary Fig. 3a). The linear regression model constructed to describe oocyst permeability to CPA attributes the observed variability to oocyst age and indicates that CPA toxicity can be manipulated by application of an appropriate bleaching protocol and CPA incubation time (*R*^2^=0.62, *p*<0.0001) (Supplementary Table 1).

### Ultra-fast cooling as an approach to oocyst cryopreservation

Because the CPA concentration in permeabilized oocysts is not known, ultra-fast cooling rates were used as vitrification can occur with reduced CPA concentrations. Ultra-fast cooling was achieved by loading oocysts into 200 µm diameter silica microcapillaries followed by rapid submersion in liquid nitrogen (Fig. 2a). As a consequence of the small microcapillary size, the reduced thermal mass of the sample allows for a rapid cooling rate of ∼4000 K/s, such that vitrification of the CPA cocktail is nearly instantaneous (*11, 19*). Based on the DMSO toxicity data for permeabilized oocysts (Fig. 1d), a CPA cocktail solution of 30% DMSO and 0.5 M trehalose was chosen for cryopreservation experiments. This concentration ensures that vitrification will occur outside the oocyst, yet exhibits minimal toxicity following incubation for up to 20 min. While higher toxicity rates were observed in older oocysts, toxicity could be minimized by altering the bleaching protocol (Supplementary Figs. 3a and 3b).

**Fig 2:**
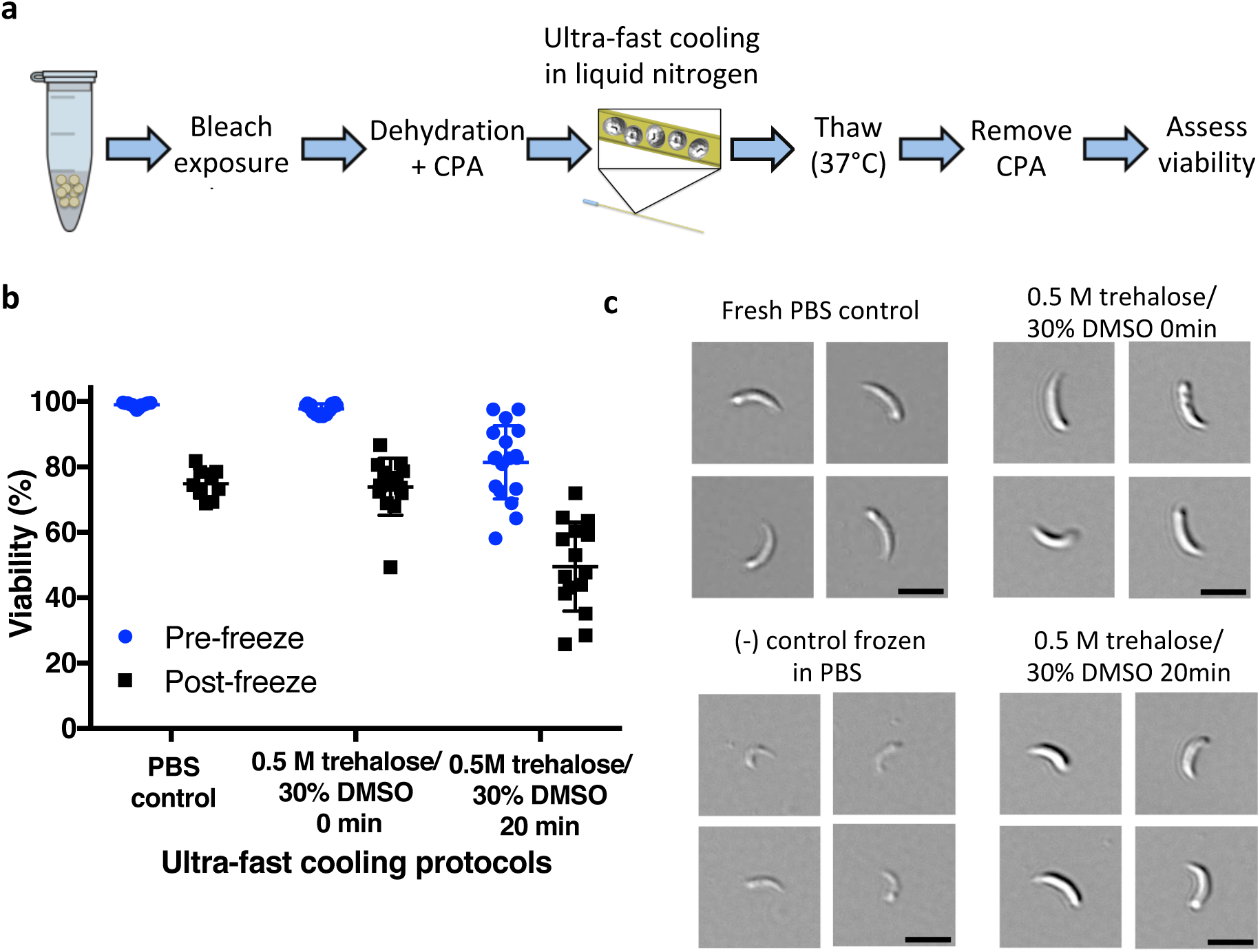
Vitrified *C. parvum* oocysts are viable and excyst. **a** To ensure vitrification, bleached oocysts were dehydrated in 1 M trehalose and then suspended in 0.5 M trehalose/30% DMSO. Oocysts were then immediately loaded into microcapillaries and plunged into liquid nitrogen for 5 min, or incubated with CPA solution for 20 min followed by loading of microcapillaries and freezing. Oocysts were thawed by quickly transferring the microcapillary from liquid nitrogen to a 37°C water bath. Viability was quantified after CPA removal based on PI exclusion and quality of excysted sporozoites. **b** Oocyst viability was determined by PI exclusion, both pre-freeze and after cryogenic storage (n=12). Values indicate means and error bars indicate standard deviation. **c** DIC micrographs show excysted sporozoites. Sporozoite viability was assessed on basis of their shape, structure and observed motility, after excystation in 0.75% taurocholic acid at 37°C. Scale indicates 5 µm.

Figure 2a shows the workflow used in ultra-fast cooling experiments. Oocysts permeabilized by hypochlorite treatment were suspended in 1 M trehalose to promote dehydration, followed by the addition of DMSO to achieve a final concentration of 0.5 M trehalose/30% DMSO. Samples were then loaded into microcapillaries and either immediately submerged in liquid nitrogen (0 min) or incubated with the CPA for 20 min prior to submersion. PBS only (no CPA) oocysts were selected as control. Figure 2b shows the pre-freeze toxicity associated with CPA exposure and post-thaw toxicity caused cumulatively by CPA exposure and cryopreservation, where the latter group exhibited increased toxicity. Cryopreservation using protocols with 0 min and 20 min DMSO incubation yielded, respectively, 73.9±8.7% and 49.5±13.6% viable oocysts, based on propidium iodide (PI) exclusion. Interestingly, 74.9±4.4% of the control oocysts frozen in PBS were also PI^-^ upon thawing. Thus, for better *in vitro* assessment of oocyst function, a PI exclusion assay was coupled with an *in vitro* excystation assay to assess the ability of oocysts to release viable sporozoites. The viability of excysted sporozoites was assessed on the basis of shape, structure and observed motility (Fig. 2c). Despite the high viability rates inferred from PI exclusion, sporozoites excysted from oocysts frozen in PBS were morphologically damaged and non-infectious in subsequent animal experiments (Supplementary Fig. 4). Oocysts frozen after either 0 min or 20 min CPA exposure yielded morphologically normal, full-bodied and often motile sporozoites, which resembled those excysted from unfrozen control oocysts. Based on the promising results obtained from this study, both the 0 and 20 min incubation protocols were selected for subsequent experiments to assess infectivity of thawed oocysts in mice.

### Oocyst infectivity in the IFN-γ knockout mouse model

The infectivity of cryopreserved oocysts was tested in the IFN-γ knockout mouse model. Because the volume of the oocyst suspension was limited by the size of the microcapillaries, the choice of animal model was dictated by its high susceptibility to infection with a low infectious dose of *C. parvum* oocysts (*20*). Further, because oocyst age was found to play a significant role in CPA toxicity (Supplementary Fig. 3, Supplementary Table 1), cryopreservation outcome was measured among a single batch of oocysts aged 1, 6 or 12 weeks. Figures 3a-c show that oocysts cryopreserved by ultra-fast cooling using either the 0 min or 20 min incubation protocol were infectious to mice. Specifically, each mouse inoculated with cryopreserved oocysts developed a patent infection at 5-6 days post infection (DPI), similarly to unfrozen control oocysts, as evidenced by onset of fecal oocyst shedding. Histology of intestinal tissue demonstrates the extent of the infection in all treatment groups, except for mice infected with heat-inactivated control oocysts (Fig. 3d). The micrographs of jejunal sections demonstrate heavy infestation of epithelial cells with the parasite. Oocyst shedding was delayed in mice infected with 6- and 12-week-old oocysts in comparison to 1-week-old oocysts, indicating that the infectivity of oocysts was lower for older inocula treated with either cryopreservation protocol. Interestingly, this data demonstrates that oocysts younger than 12 weeks can be successfully cryopreserved using either 0 min or 20 min protocols. Alternatively, 16-week-old oocysts can only be cryopreserved using the 20 min treatment with DMSO following trehalose dehydration (Supplementary Fig. 5).

**Fig 3:**
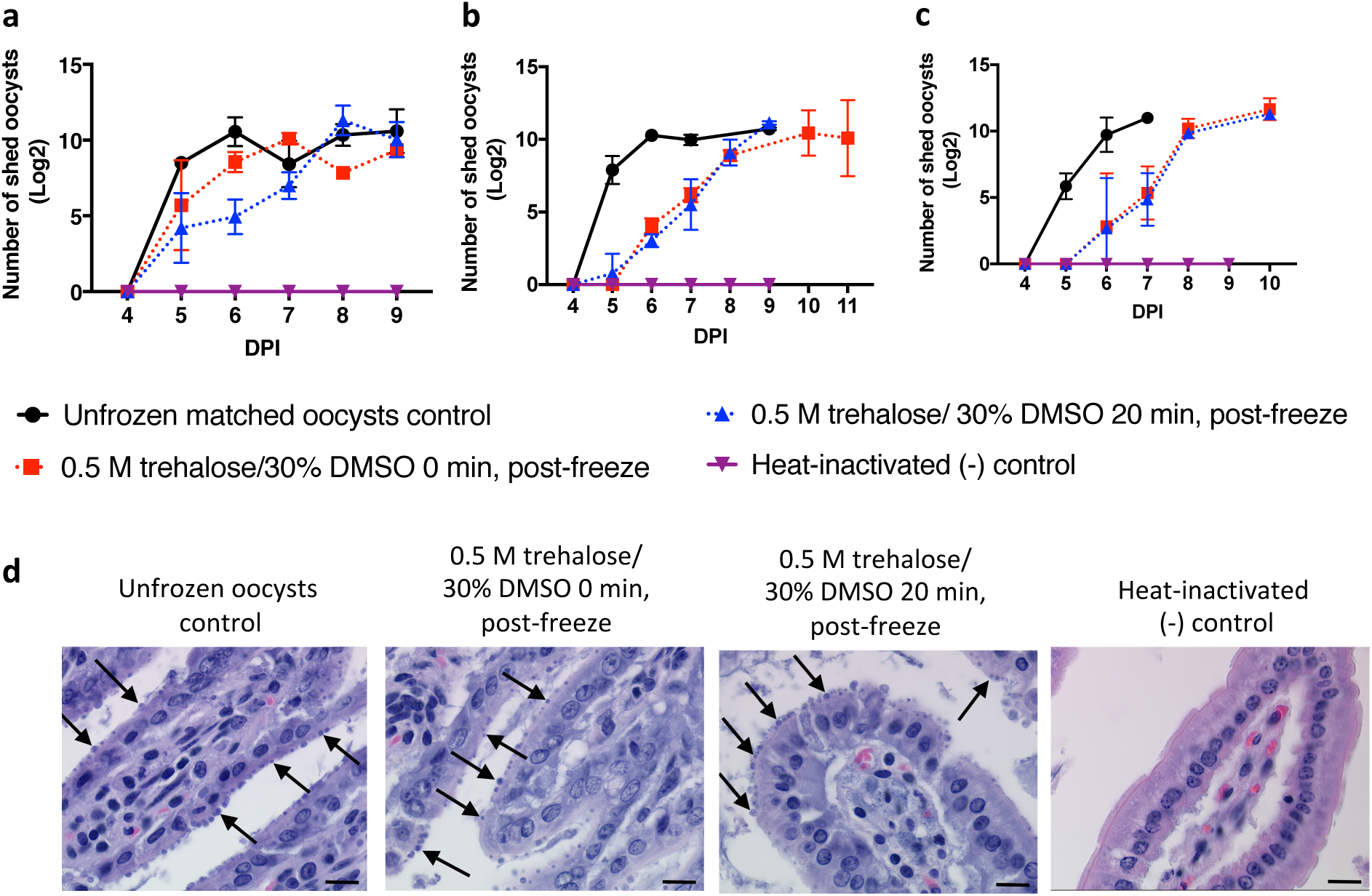
Cryopreserved *C. parvum* oocysts are infectious to IFN-γ knockout mice. Dehydrated oocysts were vitrified using either the 0 min or 20 min DMSO incubation protocol. IFN-γ knockout mice were inoculated orally with 5,000 PI^-^ thawed or unfrozen control oocysts. Intensity of fecal shedding was quantified daily by microscopic enumeration of oocysts in 30 fields of acid-fast stained fecal smears under 1000x magnification. Positive (unfrozen) and negative controls (heat-inactivated) were included as matched controls. To determine whether oocysts age has effect on the success of the cryopreservation protocol, **a** 1-**b** 6- and **c** 12-week-old oocysts were studied (n=2-3 mice). Values indicate means of log transformed oocysts count and error bars indicate standard deviation. Viability of oocysts was determined by PI exclusion prior to inoculation and was as follows: a) 43.1% and 76.7% b) 45.2% and 80.6% c) 53.1% and 72.4%, for 20 min and 0 min incubations, respectively. **d** Micrographs of hematoxylin and eosin–stained jejunal sections from IFN-γ mice infected with fresh, cryopreserved and heat-inactivated 12-week-old oocysts. Arrows indicate intracellular stages of the parasite located in the apical region of intestinal epithelial cells. Scale indicates 20 µm. Data regarding infectivity of each individual mouse can be found in Supplementary Fig. 6.

While this data suggests that 20 min is sufficient to permit intracellular accumulation of DMSO (Fig.1d, Fig. 4, Supplementary Fig. 2), it is unlikely that any substantial DMSO accumulation occurs using the 0 min protocol, though substantial dehydration is observed (Fig. 4). The observation that 16-week old oocysts require a 20 min incubation with the CPA provides some explanation of the phenomenon of successful cryopreservation with 0 min protocol in younger oocysts. We suspect an age dependent depletion of naturally present biomolecules, which can act as vitrifying agents, occurs over the course of 16 weeks. While the intracellular uptake of DMSO may vary dependent upon oocyst age, we found that the presence of both trehalose and DMSO is essential for successful cryopreservation using the ultra-fast cooling method (Supplementary Fig. 4).

**Fig 4:**
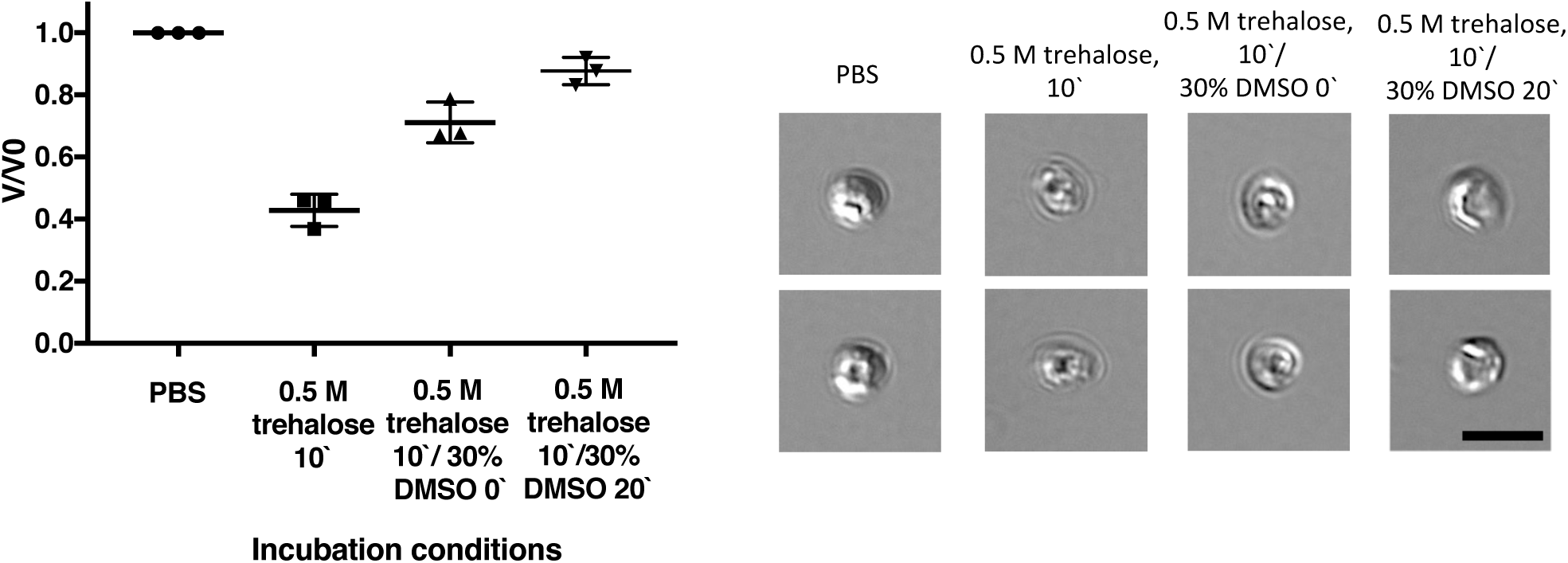
Response of *C. parvum* to dehydration and CPA uptake prior to freezing. The shrink-swell response of bleached oocysts to a challenge with 1 M trehalose for 10 min, with or without subsequent addition of DMSO for 0 min or 20 min (at final concentration of 0.5 M trehalose/ 30%DMSO) was measured by image analysis and compared to the PBS control. Loss of 57.2 ±5.2% of oocyst volume related to dehydration in trehalose is partly restored by addition of DMSO. This is likely related to a decreased trehalose gradient after immediate addition of DMSO (0 min) or intra-oocyst uptake of DMSO during 20 min incubation (n=3). Values indicate means and error bars indicate standard deviation. DIC micrographs demonstrate volumetric changes of oocysts. Scale indicates 5 µm.

To determine whether the oocysts remain stable during storage, we cryopreserved a single batch of oocysts using the ultra-fast cooling protocol with 0 min exposure to DMSO and measured infectivity after 1, 4 or 12 weeks of cryogenic storage (Fig. 5). No impact on cryopreservation outcome was observed as a consequence of storage duration. We further tested this protocol using 1.5 mL Eppendorf tubes with higher sample volumes and a slow cooling rate of 1°C/min. However, this approach did not yield infectious oocysts (Supplementary Fig. 4), confirming that rapid cooling rates are essential.

**Fig 5:**
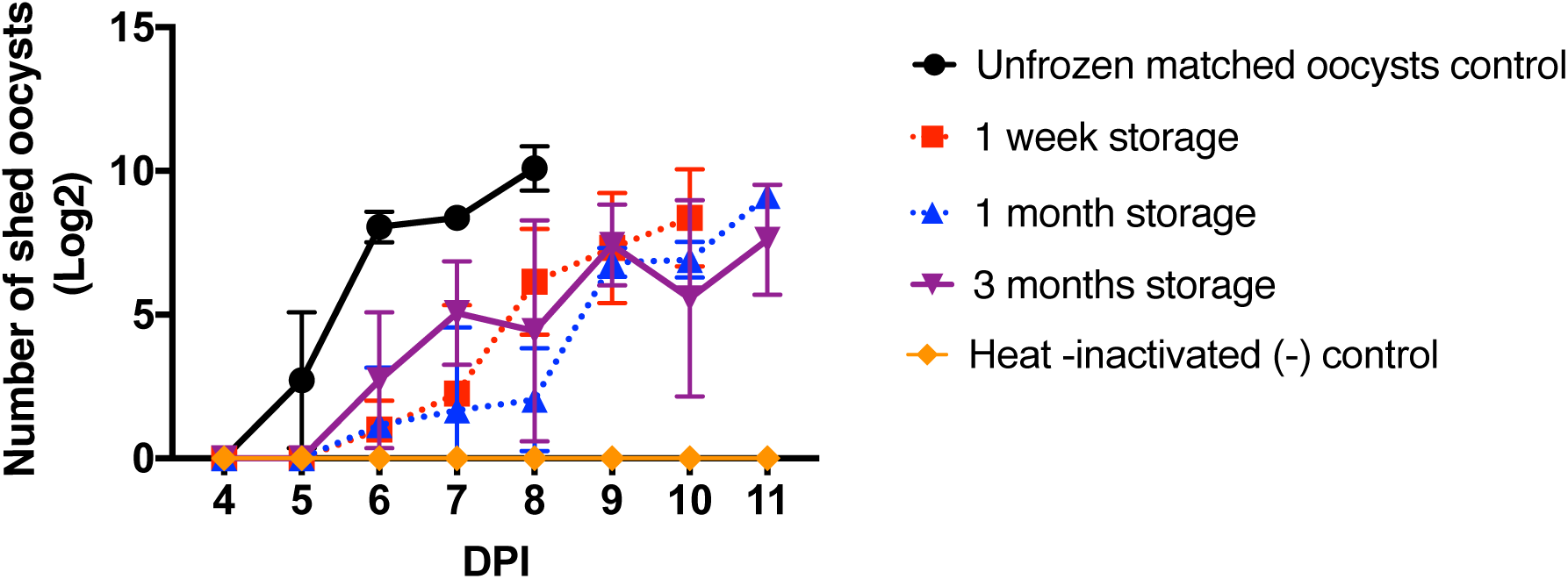
Long-term cryogenic storage does not decrease infectivity of cryopreserved oocysts. Bleached *C. parvum* oocysts (5%, 1 min) were frozen using the ultra-fast cooling protocol following dehydration in trehalose (10 min) and 0 min incubation in 0.5 M trehalose/30% DMSO solution. Oocysts were stored in microcapillaries in liquid nitrogen and thawed after 1, 4 and 12 weeks. Positive (unfrozen) and negative controls (heat-inactivated) were included as matched controls. IFN-γ knockout mice were inoculated orally with 5,000 PI^-^ oocysts (n=3). Intensity of fecal shedding was quantified daily by microscopic enumeration of oocysts in 30 fields of acid-fast stained fecal smears under 1000x magnification. Values indicate means of log transformed oocysts count and error bars indicate standard deviation. Mice inoculated with cryopreserved oocysts developed similar infection at 6dpi regardless of the storage length, as evidenced by fecal shedding of oocysts. Viability of oocysts was determined by PI exclusion prior to inoculation and was as follows: 82.6%, 79.1% and 80% for 1, 4 and 12 weeks of storage respectively. Data regarding the infectivity of each individual mouse can be found in Supplementary Figure 7.

## Discussion

This is the first report of a successful cryopreservation protocol for infectious *Cryptosporidium* oocysts. Cryopreservation was achieved by incubation of oocysts in a CPA cocktail of trehalose and DMSO, followed by ultra-fast cooling. The presence of both trehalose and DMSO coupled with rapid cooling rates was essential for successful cryopreservation. While this CPA cocktail is sufficient to vitrify with the cooling rates used in this study, it is unclear whether the oocyst itself actually vitrifies. Interestingly, both 0 and 20 min exposures to the CPA cocktail resulted in successful cryopreservation in oocysts aged ≤12 weeks. While a 20 min incubation is sufficient to permit CPA uptake, the 0 min incubation likely results in little or no CPA accumulation, suggesting intra-oocyst CPA may not be necessary. We consider two hypotheses to account for oocyst survival in the absence of CPA uptake. First, the presence of native sugars and other molecules in oocysts may be sufficient for vitrification (*20*). Consistent with this hypothesis is the necessity to dehydrate oocysts prior to cryopreservation, thereby concentrating intracellular contents, which would increase the potential for vitrification. Further, a decreased infectivity of aged oocysts after cryopreservation indicated by a delay in oocyst shedding in mice may be explained by a progressive depletion of intracellular nutrients as the oocysts age (*3*). This hypothesis is supported by analysis of 16-week old oocysts, where only those incubated with CPA for 20 min (sufficient time for CPA uptake) were successfully cryopreserved. An alternative hypothesis is that small ice crystals may be present within the oocysts, but at insufficient sizes to induce damage. Regardless of the mechanism by which the parasite survives cryopreservation, this method was demonstrated to preserve viable, infectious *C. parvum* oocysts and represents a significant contribution to the field.

We recognize two key limitations of this study are the limited number of strains examined and the small volume capacity of the microcapillary. While both isolates of *C. parvum* examined here were found to respond favorably to the cryopreservation method and showed no significant differences in viability and infectivity after thawing, additional strains must be evaluated to determine how widely applicable this method is. It would be especially interesting to determine whether this protocol is effective for *C. hominis* and transgenic strains of *C. parvum* (*21*). To address the volume limitations imposed by the microcapillary, we are now exploring methods to enable >100 µl storage volumes. Potential approaches include the use of commercially available insemination straws that are available in larger volumes (100-250 µl). Due to the increased thermal mass, it may be necessary to incorporate rapid rewarming methods or small molecules to overcome ice crystallization during sample thawing (*22, 23*).

The cryopreservation method reported here for *Cryptosporidium* oocysts will allow scientist to maintain unique strains of the parasite, field isolates and genetically modified laboratory strains, without the need of propagation in susceptible animals. Most importantly, cryogenic storage will enable standardized testing of therapeutic and vaccine candidates in animal experiments and human challenge studies. This is currently hindered by batch-to-batch variation in oocyst viability and infectivity. The ability to cryopreserve single batches of oocysts will not only ascertain uniformity of the infectious dose used in drug efficacy studies but will also eliminate the need for optimization, standardization and validation among multiple batches of oocysts, a cumbersome and time-consuming process.

## Methods

### Oocyst sources and collection method

*C. parvum* oocysts (Iowa isolate, isolated from infected calves) were purchased from Bunch Grass Farms (Deary, ID) and stored at 4°C in an antibiotic solution containing 100 U/ml penicillin, and 100 µg/ml streptomycin. *C. parvum* oocysts, MD isolate, were generated at the Tufts University by propagation in 8-week-old CD-1 mice (Charles River Laboratories, Wilmington, MA). Three days preceding infection, mice were placed in bedded microisolators and immunosuppressed with dexamethasone in drinking water at a concentration 16 mg/l, which continued throughout the course of experiment. Each mouse was inoculated orally with 50,000 oocysts in PBS suspension. Starting from onset of shedding on 5 DPI, mice were placed for ten consecutive days in collection microisolators for overnight collection of feces and rested in bedded microisolators during the day, both with access to food and water *ad libitum*. Mice were euthanized on 15 DPI by carbon dioxide asphyxiation, followed by cervical dislocation as required by the Tufts University IACUC guidelines. Oocysts were purified from mice feces using a multi-step protocol including ether extraction and nycodenz gradient steps, as described elsewhere (*26*) with minor modifications. Briefly, feces were homogenized and filtered through a 100 µm cell strainer. Fat was removed from the fecal slurry by treatment with diethyl ether in water containing 0.5% Tween 80 at the ratio 2:3. Oocysts were separated from the fecal material by ultracentrifugation (13,400 ×g, 60 min) on a 10%/25% nycodenz gradient. Oocysts were then resuspended in PBS and stored in 4°C.

### Oocyst bleaching

When indicated, oocysts were bleached using dilutions of commercial bleach containing 8.25% sodium hypochlorite (Clorox Original, The Clorox Company, CA). Following incubation with bleach on ice (1-7 min), oocysts were washed three times by suspension in PBS and centrifugation (18,000 ×g, 2 min). A more detailed bleaching protocol can be found in the Supplementary Information. Viability of bleached oocysts and integrity of oocysts wall was measured by exclusion of PI at 10 µg/ml and carboxyfluorescein succinimidyl ester (CFSE) at 5 µM concentration, respectively, by flow cytometry. Impact of bleaching on oocyst excystation was determined by excystation rate in response to treatment with 0.75% taurocholic acid at 37°C for 1 hour (Supplementary Fig. 1).

### Permeability of oocysts to extracellular CPAs

To determine oocyst permeability to CPAs, unbleached *C. parvum* oocysts (Iowa isolate) were incubated with 40% solutions of CPAs in PBS: DMSO, ethylene glycol or propylene glycol, for 30 min-2 h at room temperature. CPA was then diluted out by incubation in excess of PBS (at ratio 1:100) for 30 min at room temperature and removed by centrifugation (18,000 ×g, 2 min). Cytotoxicity attributed to CPA shown in Fig. 1a was measured by PI exclusion using flow cytometry and was normalized to its control in PBS for each time point. The experiment was performed in triplicate.

Figure 1b describes use of hypochlorite treatment as a method to increase permeability to CPAs. Unbleached and bleached (5% bleach, 7 min) *C. parvum* oocysts (Iowa isolate) were incubated with 40% DMSO for 30 min-2 h at room temperature. DMSO was then diluted with excess PBS (1:100) and cytotoxicity attributable to DMSO was measured by PI exclusion. The experiments were performed in triplicate.

The kinetics of cytotoxicity induced by dehydration (Figure 1d) were measured in oocysts treated with DMSO with or without prior dehydration. Bleached (5% bleach, 7 min) *C. parvum* oocysts (MD isolate) were dehydrated in 1 M trehalose or incubated in PBS for 10 min. Oocysts were then incubated in 30-40% DMSO or PBS for 5-60 min at room temperature. DMSO was then diluted in excess PBS (1:100) and cytotoxicity attributable to DMSO was measured as described above. The experiment was performed in triplicate.

### CPA toxicity as a function of oocyst age

Variability in CPA toxicity observed between oocyst stocks was measured individually as a function of DMSO incubation conditions, oocyst age, and bleaching method (Supplementary Figs. 3a and 3b) and collectively in a linear regression model (Supplementary Table 1). Briefly, *C. parvum* oocysts (MD isolate) at ages of 1, 4, 6, 8 and 12 weeks, were bleached using a concentration of 20%, 10% or 5% bleach (1 min), dehydrated in 1 M trehalose (10 min) and then incubated with DMSO to achieve a final concentration of 30% DMSO/0.5 M trehalose solution. Oocysts were then incubated for 10-30 min at room temperature. DMSO was then diluted with excess PBS (1:100) and cytotoxicity attributable to DMSO was measured as described above. Data was collected in triplicate and used to construct a linear regression model, in which three independent variables: oocysts age*, bleach concentration** and DMSO incubation time***, describe the outcome variable- cytotoxicity attributable to DMSO. The dataset has met the following requirements for the inclusion in a linear regression model. Normality of outcome variable was met by invocation of the central limit theorem (n=236). Each independent variable demonstrated a required linear relationship with the outcome variable (p=0.012*, p<0.0001**, p<0.0001***). The model was ran as robust due to failure of one variable to meet the requirement of constant variance (Breusch-Pagan test; p=0.01***).

### Evaluation of oocyst dehydration in hyperosmotic solutions

*C. parvum* oocysts (Iowa isolate) were incubated at room temperature for 10 min with NaCl solution at concentrations ranging from 165-500 mM or trehalose from 330-1000 mM, in addition to control oocysts incubated in PBS. Oocyst volumes shown in Fig. 1c were measured by flow cytometry (Accuri C6, BD Life Sciences) and regressed from the forward scatter signal (linear relationship between particle diameter and forward scatter at *R*^2^=0.93). Micrographs (400x) of oocysts were taken using Nikon Eclipse Ti-E microscope (Nikon Instruments Inc.). The experiment was performed in triplicate. One-way ANOVA was performed to determine significant differences between volume of untreated and treated oocysts.

### Shrink and swell response of oocysts in presence of DMSO

Supplementary Fig. 2 demonstrates the timeline of DMSO uptake by bleached oocysts. *C. parvum* oocysts (Iowa isolate) were pre-treated with bleach (5%, 7 min), washed in PBS and then incubated with 30% DMSO for 15 min in room temperature. Micrographs of oocysts were taken every minute of incubation under 400x magnification (Nikon Eclipse Ti-E microscope). Oocyst volume was measured using NIS Elements image analysis software (Nikon Instruments Inc., Melville, NY). Segmentation of oocysts was defined by application of threshold for circularity at 0.5-1 and diameter at 1-10 µm. Volume was calculated from the averaged oocyst area. The experiment was performed in triplicate.

CPA uptake by dehydrated oocysts after 0 and 20 min incubation with DMSO is shown in Figure 2. Bleached (5% bleach, 7 min) and dehydrated (1 M trehalose, 10 min) *C. parvum* oocysts (MD isolate) were incubated with DMSO (30% DMSO/0.5 M trehalose final concentration) for 20 min in room temperature. 400x and 600x (DIC) micrographs of oocysts were taken using Nikon Eclipse Ti-E microscope (Nikon Instruments Inc.) before and after DMSO treatment at 0 min and 20 min incubation time points, as well as control oocysts suspended in PBS. Diameters of fifty oocysts were measured individually for each condition using NIS Elements image analysis software (Nikon Instruments Inc., Melville, NY) and used to calculate volume. The experiment was performed in triplicate.

#### Microcapillaries

Fused silica microcapillaries with an inner diameter of 200 µm (item number Z-FSS-200280) were purchased from Postnova Analytics Inc. (UT, USA) and cut into 7.5 cm lengths. Tygon tubing was then put on the end of the microcapillary to enable use of a needle to dispel contents. Capillary action was used to load oocysts.

#### Cryopreservation protocol for ultra-rapid cooling of oocysts

Figure 2a demonstrates the workflow used in cryopreservation experiments. It is critical to follow the protocol closely, as deviations may lead to lethal ice crystallization. A detailed step-by-step protocol, based on the “0” min experiment, is included in the Supplementary Information (Supplementary Protocol 1). Briefly, 1,000,000 *C. parvum* (MD and Iowa isolates) oocysts originating from the same stock were bleached as follows: ≤4-week-old oocysts with 20% bleach (1 min), ≥6-week-old oocysts with 5% bleach (1 min). Oocysts were centrifuged (18,000 ×g, 2 min), resuspended in 1 M trehalose and incubated in room temperature for 10 min. 60% DMSO was added to dehydrated oocysts to achieve the final concentration of 0.5 M trehalose/30% DMSO and incubated for “0” or 20 min in room temperature. Following incubation with CPA, oocysts were loaded into microcapillaries. Three microcapillaries per condition were loaded by capillary action with approximately 400,000 oocysts and then plunged into liquid nitrogen without sealing. For all “0” min experiments, except for long-term storage, the exposure to CPA during loading was <1 min prior to freezing. Though the sample reaches cryogenic temperatures nearly instantaneously, oocysts were maintained in liquid nitrogen for 5 min prior to thawing. Thawing was achieved by quickly transferring the microcapillary from liquid nitrogen to a 37°C water bath. It is critical that the microcapillary be rapidly transferred into liquid nitrogen, as well as be rapidly transferred from liquid nitrogen to 37°C for thawing, in order to avoid lethal ice crystallization. Microcapillaries were expelled using a 1 ml syringe and a PrecisionGlide needle 30 ga. (BD), and incubated in 1 ml of PBS in room temperature for 30 min to relieve the cryogenic stress. The remainder of unfrozen oocysts was also resuspended in 1ml PBS to assess pre-freeze cytotoxicity. PBS was then removed from both frozen and unfrozen samples by centrifugation (18,000 ×g, 2 min) and 100 µl of PBS was used to resuspend the pellet. Thawed oocysts were stored on ice prior to further testing.

Supplementary Fig. 4 demonstrates infectivity of oocysts cryopreserved using the protocol described above with modifications including exclusion of trehalose, exclusion of DMSO and use of a slow cooling rate. In the slow cooling method, oocysts were placed in 1.5 ml Eppendorf tubes instead of microcapillaries and frozen to -80°C in a Mr. Frosty freezing container (Thermo Scientific) at cooling rate 1°C/min.

#### *In vitro* viability of cryopreserved oocysts

Bleached *C. parvum* oocysts (MD isolate) were cryopreserved using 0.5 M trehalose/30% DMSO with a 0 min or 20 min CPA incubation period, as described above. Control oocysts were frozen in PBS in an identical manner. Pre-freeze and post-freeze oocysts viability, shown in Fig. 2b, was measured by propidium iodide staining at 10 µg/ml using flow cytometry. As we found inclusion of PI alone to be an inaccurate marker of viability, consistent with reports from other groups (*24, 25*), we coupled oocyst viability with rate of excystation and quality of excysted sporozoites. To achieve excystation, oocysts were incubated with 0.75% taurocholic acid at 37°C for one hour. The excystation rate was calculated as a percentage of excysted oocysts. Quality of excysted sporozoites was assessed by microscopic inspection on basis of their shape, structure and observed motility. Data was collected in triplicate for each oocysts age tested (total n=12). DIC micrographs (600x) of sporozoites shown in Fig. 2c, were taken using Nikon Eclipse Ti-E microscope (Nikon Instruments Inc.).

#### *In vivo* infectivity of cryopreserved oocysts

A summary table describing the different cryopreservation methods evaluated and the corresponding outcome can be found in Supplementary Table 2. Briefly, *C. parvum* oocysts of MD isolate at 1, 6 and 12 weeks (Fig. 3a-c) of age and of Iowa isolate at 10 and 16 weeks of age (Supplementary Fig. 5) were bleached and cryopreserved as described above using the 0 min or 20 min incubation time in 0.5 M trehalose/30% DMSO. Infectivity of cryopreserved oocysts was tested in 8-week-old female IFN-γ knockout mice (obtained Jackson Laboratory, Bar Harbor, ME, bred under pathogen-free conditions). Mice were pooled in randomly assigned groups and housed in bedded microisolators with access to food and water *ad libitum*. Mice were immunosupressed with dexamethasone in drinking water at concentration 10 mg/l. Dexamethasone treatment started on three days preceding infection and continued throughout the course of the experiment. Each mouse was inoculated orally with 5,000 PI^-^ cryopreserved oocysts in PBS suspension. For each batch of experiments, a positive control group (mice inoculated with 5,000 fresh PI- oocysts) and negative control group (mice inoculated with 5,000 heat-inactivated oocysts) were included. Mice feces were collected daily from each animal individually starting from 4 DPI. Mice were euthanized by carbon dioxide asphyxiation, followed by cervical translocation on 11 DPI unless prematurely terminated and excluded from the analysis due to recumbency. Intensity of fecal shedding was quantified daily by inspection of the acid-fast stained fecal smear slides for presence of oocysts in 30 fields (1000x). Small and large intestines were collected from each mouse *post mortem* for preparation of histological sections stained with hematoxylin and eosin. Micrographs of histological sections (500x) shown in Fig. 3d, were taken using Olympus BX41TF microscope (Olympus, Tokyo, Japan). The investigators were not blinded during this experiment.

#### Long-term storage of oocysts in liquid nitrogen

The cryopreservation protocol is sensitive to variations in technique. We have included a more detailed description of the protocol in the Supplementary Information. Briefly, one million oocysts of 10-week-old *C. parvum* (Iowa isolate) were treated with 5% bleach (1 min) and cryopreserved as described above using the ultra-fast cooling protocol following dehydration with trehalose (1 M) and addition of DMSO (final concentration 0.5 M trehalose and 30% DMSO). Oocysts were immediately loaded into ten microcapillaries and plunged into liquid nitrogen to minimize uptake of DMSO, referred to as 0 min incubation. While the 0 min incubation period actually ranged from 45 s – 3 min of exposure to CPA prior to submersion in liquid nitrogen, it is unlikely any substantial intracellular CPA accumulation occurred, as indicated by the shrink swell response of oocysts in DMSO solutions (Supplemental Figure 2). Afterwards, microcapillaries were transferred into 5 ml cryovials (Globe Scientific) with perforated caps, which were fixed to cryocanes (Thermo Fisher Scientific) and placed in ladles of the nitrogen tank for liquid phase storage (Bio-cane, Thermo Fisher Scientific) for up to 12 weeks. Microcapillary transfer into cryocanes was performed in a Styrofoam box to ensure that the microcapillary remains submerged in liquid nitrogen at all times. Thawing was performed as described above. Infectivity of cryopreserved oocysts was tested in 8-week-old IFN-γ knockout mice (Jackson Laboratory, Bar Harbor, ME) as described above and is shown in Figure 5.

#### Statistical analysis

All the *in vitro* experiments were performed in three repeats and replicated three times to ensure reproducibility. All the *in vivo* experiments were performed with 3 animals in each group, which is the minimum amount of animals to test for oocyst infectivity. Graphing and statistical analyses of data were done using GraphPad Prism software (v7, GraphPad Software, Inc.) and Stata 15 (StataCorp. 2017).

## Acknowledgments

The research was supported by the BMGF grant OPP1139037.

## Data availability

The datasets generated during and/or analyzed during the current study are available from the corresponding author on reasonable request.

## Ethical compliance

Animal experiments were conducted at the Tufts University in compliance with study protocols approved by the Institutional Animal Care Use Committee.

## Author contributions

J.J. and R.S. contributed equally. All authors discussed the results, data analysis and the paper.

